# Reanalysis of double EphA3 knockin maps in mouse suggests that stochasticity in topographic map formation acts within the retinal cell population rather than between competing mechanisms at the colliculus

**DOI:** 10.1101/2022.03.29.486226

**Authors:** David J Willshaw, Nicholas M Gale

## Abstract

It has been suggested that stochasticity can act in the formation of topographically ordered maps in the visual system through the opposing chemoaffinity and neural activity forces acting on the innervating nerve fibres being held in an unstable equilibrium. Evidence comes from the Islet2-EphA3 knockin mouse, in which approximately 50% of the retinal ganglion cells, distributed across the retina, acquire the EphA3 receptor, thus having an enhanced density of EphA which specifies retinotopic order along one axis of the retinocollicular map. Sampling EphA3 knockin maps in heterozygotes at different positions along the mediolateral extent of the colliculus had found single 1D maps (as in wild-types), double maps (as in homozygous knockins) or both single and double maps. We constructed full 2D maps from the same dataset. We found either single maps or maps where the visual field projects rostrally, with a part-projection more caudally to form a double map, the extent and location of this duplication varying considerably. Contrary to previous analyses, there was no strict demarcation between heterozygous and homozygous maps. These maps were replicated in a computational model where, as the level of EphA3 was increased, there was a smooth transition from single to double maps. Our results suggest that the diversity in these retinotopic maps has its origin in a variability over the retina in the effective amount of EphA3, such as through variability in gene expression or the proportion of EphA3+ retinal ganglion cells, rather than the result of competing mechanisms acting at the colliculus.

**Significance statement:** Analysis by others of visuocollicular maps in EphA3 knockin mice indicated stochasticity in the development of nerve connections. Here a proportion of the retinal ganglion cells have the chemoaffinity ligand EphA3. Sampling the heterozygous map revealed either a single map (as in wildtypes), a double map (as in homozygotes) or a mixture, suggesting an unstable balance between competing chemoaffinity and neural activity forces. We constructed full 2D maps from the same dataset. We found a diversity of double maps in both genotypes, with no demarcation between heterozygote and homozygote, replicated in a computational model where the level of EphA3 varies between animals. We suggest that any stochasticity acts at the level of the retina rather than downstream at the colliculus.

## Introduction

In mammals, retinal ganglion cells develop an ordered map of nerve connections across the superior colliculus (Drager and Hubel, 1976; Finlay et al., 1978; King et al., 1996), driven by both chemical and electrical signalling mechanisms (Pfeiffenberger et al., 2006; Cang and Feldheim, 2013). The mapping of nasotemporal retina to rostrocaudal colliculus involves the EphA receptor tyrosine kinases and their ligands, the ephrinAs, distributed as gradients across the nasotemporal and rostrocaudal axes, respectively. The mapping of dorsoventral retina to mediolateral colliculus is mediated by gradients of EphB and ephrinB running dorsoventrally and mediolaterally (Flanagan and Vanderhaeghen, 1998; McLaughlin et al., 2003a).

In a wildtype (WT) map there is a single ordered projection of the retinal nasotemporal axis onto the collicular rostrocaudal axis (Brown et al., 2000). In an EphA3 knockin mouse, the EphA3 ligand, not normally present in mouse retina, is bound to the Islet2 class of retinal ganglion cells which are distributed over the retina in equal proportions in a salt-and-pepper fashion (Brown et al., 2000). In the EphA3^ki/ki^ map, EphA3+ retinal cells project to rostral colliculus, with a similarly ordered projection from EphA3− cells more caudally, to form a double representation of visual field on colliculus. In the EphA3^ki/+^ map, the two types of cells also project separately except that the projections coincide in far rostral colliculus (Brown et al., 2000; Reber et al., 2004; Bevins et al., 2011). It was inferred that the degree of separation of the two projections on the colliculus depends on the extra amount of EphA assigned to EphA3+ cells, in the homozygous knockins being double that in the heterozygous knockins. (Brown et al., 2000; Reber et al., 2004).

Owens et al. (2015) explored the maps in EphA3^ki/+^ animals using Fourier-based intrinsic imaging, where the visual field is scanned over two orthogonal directions. (Kalatsky and Stryker, 2003). They sampled the azimuthal scan to construct the 1D nasotemporal to rostrocaudal projection at three different dorsoventral locations. They found either single, double or a mixture of single and double maps. They proposed that this variety amongst EphA3^ki/+^ maps is evidence for stochasticity operating in the formation of nerve connections: that in the EphA3^ki/+^ animal, activity-based forces favouring single maps are pitted against chemically-based forces favouring double maps in an unstable equilibrium, small fluctuations resulting in either a single or double map or a mixture (Figures 1A-1C).

**Figure 1.**
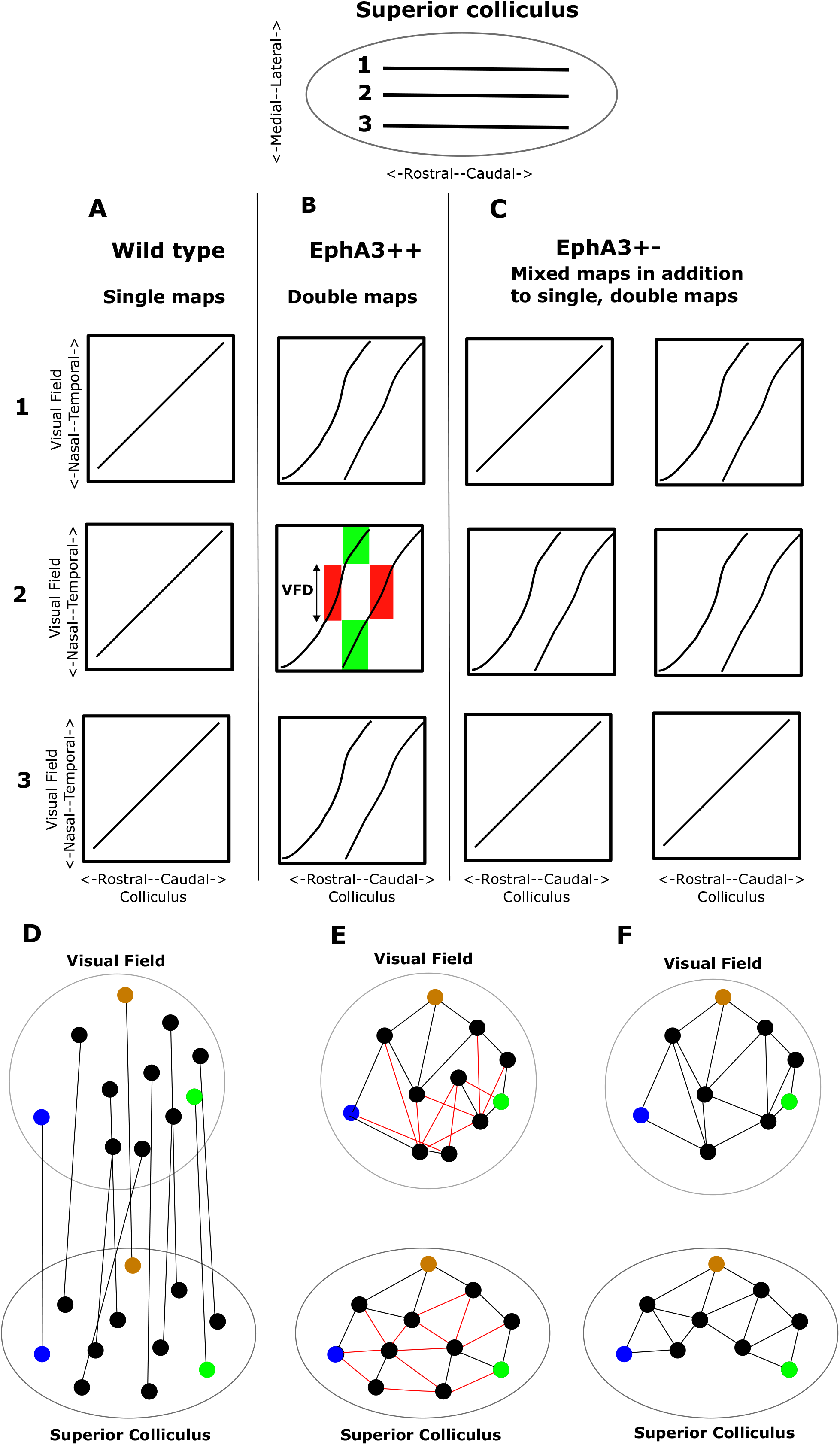
**A, B, C.** Schematic EphA3 knockin visual maps of the 1D projection along the nasotemporal axis of visual field sampled along three different tracks 1,2,3 running rostrocaudally and spaced along the lateral-medial axis of the colliculus, used to show schematically the findings of Owens et al. (2015). These maps are also used to illustrate the general properties of maps in 1D. The shapes of the 1D profiles were redrawn from Brown et al. (2000). **A.** Wild type. Mechanisms of neural activity and chemoaffinity combine to form a single projection along all three tracks. **B.** Homozygous knockin (EphA3^ki/ki^). Double maps along all three tracks. With high EphA3 in the Islet2 retinal ganglion cells, the EphA3+ and the EphA3− retinal ganglion cells each make a separate map. Green areas in panel 2 enclose the connections made from two distinct parts of the visual field to the same part of the colliculus. Red areas enclose the connections made from the same part of the visual field to two distinct regions of the colliculus, with its extent measured by the amount of Visual Field Duplication (VFD). **C**. Heterozygous knockin (EphA3^ki/+^). In addition to entirely single and entirely double maps, in some cases there is a mixture of these two types. Two possible explanations are: **The collicular hypothesis.** With a low level of EphA3 in the heterozygote, the mechanisms of neural activity and chemoaffinity on the colliculus are finely opposed in an unstable equilibrium so that all combinations of maps may be seen along the three tracks (Owens et al., 2015). **The retinal hypothesis.** Since double maps are associated with high EphA3 and single maps with zero EphA3, there is a variation of the level of EphA3 across the retina leading to different types of map along different tracks on the colliculus. **D, E, F.** Use of the Lattice Method to construct an ordered one-to-one 2D map. **D**. Virtual electrodes are placed on the colliculus to cover the surface. They are added one by one at random positions subject to the constraint that each new electrode is at a pre-specified distance Δ (+/‒10%) from all existing electrodes. The procedure terminates once no more suitable positions can be found. For each electrode the corresponding position in the visual field is then found by averaging over all visual field positions associated with the points on the colliculus within a distance Δ/2 of the given electrode position. **E**. Nearest neighbouring electrodes on the colliculus are interconnected using Delaunay triangulation to form a lattice. Each pair of visual field points (nodes) where the corresponding electrodes are nearest neighbours are also interconnected. If the map has perfect order, the collicular lattice will be replicated on the visual field. If this is not the case, links in the visual field will cross over showing that neighbourhood relations are violated. These links are coloured red and the corresponding links on the collicular lattice are also coloured red. **F**. Pairs of electrodes and the corresponding visual field nodes are removed one by one to eliminate the cross-overs from the visual field representation. Ultimately the visual field representation becomes a lattice, indicating a perfectly ordered but reduced map. The elimination method yields a lattice containing no more than 8% of the optimal number of nodes found by linear programming where different part of the map may be disconnected (Willshaw et al., 2014). Three pairs of matching coloured points in the two structures indicate the global orientation of the map. In this schematic example, two of the eleven points had to be removed, showing that the ordered map is restricted to one part of the colliculus. Various metrics can be derived from these plots, such as Map Quality (the percentage number of nodes in the perfectly ordered lattice) compared to the original as a measure of map quality, and the polarity and the magnification of the map along any specified axis.

Since the level of EphA3 assigned to Islet2 retinal ganglion cells determines whether a double or single map results, another explanation of these results is that the level of EphA3 in EphA3^ki/+^ animals varies over the retina and thus over the retinal fibres innervating different parts of the colliculus. To distinguish both hypotheses (Legend to Figure 1) requires information about the entire map from both directions of scan. We analysed the same dataset. Following Cang et al. (2008), we excluded noisy data, and also data from collicular regions where the map is discontinuous suggesting two or more visual areas mapping to the same collicular area. These cannot be discriminated since the imaging method gives a single point in the visual field for each pixel in the collicular map. We then reconstructed the 2D visuocollicular map employing the Lattice Method (Willshaw et al., 2014), finding either single maps or maps in which the visual field projected rostrally and caudally to form a double map, the extent and location of the caudal part varying considerably; there was no strict demarcation between EphA3^ki/+-^ and EphA3^ki/ki^ maps. In these maps there is no information about how the two populations of EphA3− and EphA3+ retinal ganglion cells project onto the colliculus. Therefore, to help examine possible modes of projection we used simulation results from computer models (Triplett et al., 2011; Hjorth et al., 2015) where such information is available. We found no evidence for stochasticity operating at the colliculus but instead our results suggest that upstream variabilities in EphA3 signalling may be responsible.

## Methods

### Filtering the imaging data

In Fourier-based intrinsic imaging, the visual field is scanned in two orthogonal directions to determine separately the mapping of visual field positions along the nasotemporal axis onto the colliculus (azimuthal scan) and that of positions along the dorsoventral axis of visual field (elevational scan). Following Kalatsky and Stryker (2003), we defined the location of the visual projection on the colliculus as the region where excitation from visual field in the elevational map (dorsoventral scan) was highest, and an ellipse was drawn to enclose this region. Here, the strength of the signal varied considerably and information from only the 10000 most active pixels was considered (Cang et al., 2008). By combining data from the azimuthal and elevational scans of the visual field, the visual field position providing maximal excitation at each pixel in the collicular image was then calculated (Willshaw et al., 2014). A second stage of filtering was then necessary to remove unreliable information resulting from use of the Fourier imaging method. As shown for 1D maps in Figure 1, this method yields a single position in the visual field for each position on the colliculus. This is appropriate for a normal, single projection (Figure 1A), or for the part of a double map where there is an ordered map to two separate areas of the colliculus (shown for a 1D map by the red areas in Figure 1B). In these cases, a unique position in the visual field can be identified with each point on the colliculus. The visual field position identified with each small area of colliculus varies smoothly over the colliculus. However, in cases where two small regions of visual field map to the same area of colliculus (green area, Figure 1B), this average visual field position recorded at any pixel will be very different from the two visual field positions and therefore inaccurate. This average is liable to fluctuate from pixel to pixel (Willshaw et al., 2014) and therefore pixels for which the standard deviation of the mean fluctuation was greater than three times than that in WTs were removed from the analysis.

The active pixels remaining after these two stages of filtering are called *eligible* and those discarded are called *rejected*. Visual field information derived from the set of eligible pixels was then used to construct 1D and 2D maps.

### Extension of 1D analysis

To establish a baseline for comparing directly with the method of Owens et al. (2015), we plotted out the variation of azimuthal phase, signalling the nasotemporal component of visual field, along three straight lines drawn on the colliculus, which was intended to represent the visual field at the dorsoventral elevations of −25°, 0° and +25°. We found the projection onto the colliculus from all visual field positions at the given elevation and then made a straight line fit to these points. This line was then projected back onto the visual field and the corresponding visual field positions plotted.

### Construction of 2D maps – the Lattice Method

We used the Lattice Method (Willshaw et al., 2014), developed to analyse another set of Fourier-based intrinsic imaging data of the mouse visuocollicular projection (Cang et al., 2008). Eligible pixels were chosen at random to be ‘virtual electrodes,’ subject to there being an approximate spacing of 6 pixel-widths (here equivalent to 80µm) between nearest neighbours. For each virtual electrode, we then calculated the mean position of all visual field points having corresponding collicular positions within a circle of radius 3 pixel-widths of the virtual electrode. This gave a one-to-one mapping between 150-200 virtual electrodes on the colliculus (*collicular node*s), spaced at approximately 80µm, and the same number of averaged visual field positions (*visual nodes*). The spacing between collicular nodes was chosen to be the smallest that gives highly ordered 2D maps in WTs, as found in the previous analysis (Willshaw, et al., 2014). Figure 1D shows a simple example of a one-to-one mapping.

Neighbouring collicular nodes were then interconnected to form a lattice. The nodes in the visual field which are associated with interconnected collicular nodes were also interconnected (Figure 1E). If the resulting structure constructed in the visual field is also a lattice, with no edges crossing, then local order is preserved because each pair of neighbouring visual nodes then connects to a pair of neighbouring collicular nodes. Otherwise, removal of selected pairs of matching visual and collicular nodes yields the largest ordered submap in which the remaining visual nodes form a lattice (Figure 1F). Map Quality, the proportion of nodes remaining in the largest ordered submap, is a measure of the degree of local order.

In cases where two projection areas on the colliculus were defined by the distribution of eligible pixels, two separate partmaps were also constructed from the two lattices made from the nodes in the corresponding areas. This removed the interference caused by portions of the visual field being represented in both areas of the colliculus (green areas, Figure 1B). Global properties of the map were measured in the largest ordered submap, by comparing the properties of the lattice and the corresponding visual field lattice so formed. They are assessed along two orthogonal axes after rotating the projection in visual space by 20° to allow for the fact that in WT maps the nasotemporal axis projects at this angle onto the rostrocaudal axis (Dräger and Hubel 1976; Willshaw et al., 2014).

Order along selected axes of the map was measured. Rostrocaudal polarity (RCP) is the proportion of edges in the visual lattice which project onto the colliculus in the same relative rostrocaudal order as they would in a perfectly ordered map. A score of 100% represents perfect ordering along the rostrocaudal axis, 50% indicates random order and perfect but reversed order is 0%. Mediolateral polarity (MLP) is calculated similarly. Magnification ratio is a measure of the relative extent of the projection along a specified axis. The Azimuthal Magnification (AM) is the length of each edge in the visual lattice measured along the rotated nasotemporal axis of visual field compared to the length of the corresponding edge in the collicular lattice measured along the rostrocaudal axis; similarly for the Elevational Magnification (EM).

In cases when two partmaps were constructed by partitioning the colliculus, the extent of visual field represented in both regions of the colliculus, here called the Visual Field Duplication (VFD), was measured. In a completely double map, the VFD is the entire extent of visual field. In Figure 1B, there is a partial double map and the extent of the one-dimensional VFD is shown (red area).

### Computer simulations

A computational model (Triplett et al., 2011; Hjorth et al., 2015) for the development of a retinocollicular map, here called TK2011, was used to help to understand the data. In this model, retina and colliculus are each laid out as a 2D array of cells. The mapping is built up by iteratively creating and deleting contacts to reduce a quantity called the energy, calculated over all contacts, which reflects the influences of chemoaffinity, neural activity and competition. Each cell may have more than one contact.

Each newly created contact is chosen by sampling at random from all possible pairs of cells and similarly each possible deletion is chosen from the existing set of contacts. The chemoaffinity component of the energy is calculated from the gradients of EphA and EphB labelling the two axes of the retina and the matching gradients of ephrinA and ephrin B in the colliculus. For a retinal cell with an amount R_A_ of EphA and a collicular cell with an amount C_A_ of ephrinA, the contribution to the energy for the contact between them is the product R_A_XC_A_. This favours contacts between cells with high EphA and those with low ephrinA, and vice versa, which replicates the directions of the EphA and ephrinA gradients found experimentally (Cheng et al., 1995; Flanagan and Vanderhaeghen, 1998). In the EphB/ephrinB system, high EphB maps to high ephrinB (McLaughlin et al., 2003a) and so the contribution to the energy is −R_B_XC_B._ The contribution to the energy from electrical activity favours the development of contacts from neighbourhoods to neighbourhoods and the competitive mechanism favours the creation of synapses in high-density locations and their deletion in low-density locations.

We simulated the development of the retinocollicular map using TK2011 with the code developed by Hjorth et al. (2015). To match to the number of data points selected for analysis of the experimental data, 10000 retinal cells were distributed randomly within a circular retina subject to a minimal distance between neighbours. 10000 collicular cells were distributed in the same way, over a hemi-ellipse representing the shape of the colliculus. The shapes of the EphA profiles were taken from Reber et al. (2004), with values scaled so that in the wildtype they were between 0 and 1. We used the parameter values from Triplett et al. (2011) except that the magnitude of the activity term in the energy was scaled to ensure that in the wildtype virtual injections of 1% of the retinal area innervated 1% of the colliculus (McLaughlin et al., 2003b) and we ran each simulation for a larger number of iterations, 5X10^8^ against 1.4X10^7^. The knockin of EphA3 was modelled by adding an extra amount of EphA, DR, to approximately 50% of the population of retinal cell selected randomly. Choosing a roughly equal number of cells to be EphA3+ and EphA3− removes any competitive effect due to different numbers of cells. Parameter values are summarised in the legend to Figure 7. For EphA3^ki/ki^ animals the value of DR was assumed to be twice that for EphA3^ki/+^ animals (Brown et al., 2000). To investigate any variation due to the random positioning of cells, the random assignment of retinal cells to be EphA3+ and the randomness of the addition and deletion of synapses, a number of runs on a reduced 2000X2000 system was carried out for each value of DR.

To model the experimental results precisely we then made a functional map by taking the modelled anatomical data obtained from the initial simulations and then replicating the Fourier imaging method. We simulated the repeated scanning of the retina in two orthogonal directions using a bar with a width of 2% of the linear extent of the retina (Kalatsky and Stryker, 2003). Retinal and collicular firing rates were calculated using a Poisson model, with retinal firing rate proportional to the location of the bar and collicular firing rate calculated from the transmission of retinal spikes through the contacts formed and the transmission of collicular spikes through local excitation and inhibition (Phongphanphanee, 2012). In order to extract the mapping of collicular position onto retinal position, we then carried out Fourier analysis on the time course of the activity pattern at the repetition frequency. We constructed the map from colliculus to retina, which corresponds to the experimental maps found by Fourier intrinsic imaging. We also constructed simulated maps to the colliculus from the whole retina and from the separate subpopulations of EphA3+ and EphA3− retinal ganglion cells. In the latter two cases, the scan was restricted to the subpopulation in question. All 2D maps were constructed using the Lattice Method (Willshaw et al., 2014).

To compare with the modelling results obtained by Owens et al. (2015), we also simulated an earlier version (Tsigankov and Koulakov, 2006, 2010) of this model, here called TK2006, The crucial difference between the two models is that in TK2006, each collicular cell makes a single contact. This removes the need for a competition term in the energy function but at the expense of introducing discontinuities in the mapping.

### Access to resources

For access to the experimental data, please refer to Owens et al. (2015), where the data was first reported. The simulation code for TK2011 developed by Hjorth et al. (2015) is available at GitHub - Hjorthmedh/RetinalMap. The code used for the analysis of the experimental and the simulation data is under development and the current version is at https://github.com/Nick-Gale/Lattice_Method_Analysis_EphA3_Mutants.

## Results

The dataset is from 24 mice, five wildtypes (WT), five homozygous EphA3 knockins (HOM) and 16 heterozygous EphA3 knockins (HET), previously analysed by Owens et al. (2015). We could not analyse the data which Owens et al.(2015) used to explore the roles of neural activity and genetic variability owing to a lack of matching azimuthal and elevational scans in the database.

### Distributions of eligible pixels

Figure 2 shows twelve examples of distributions of the grey (eligible) pixels used to construct the map after filtering to reject the green pixels.

**Figure 2.**
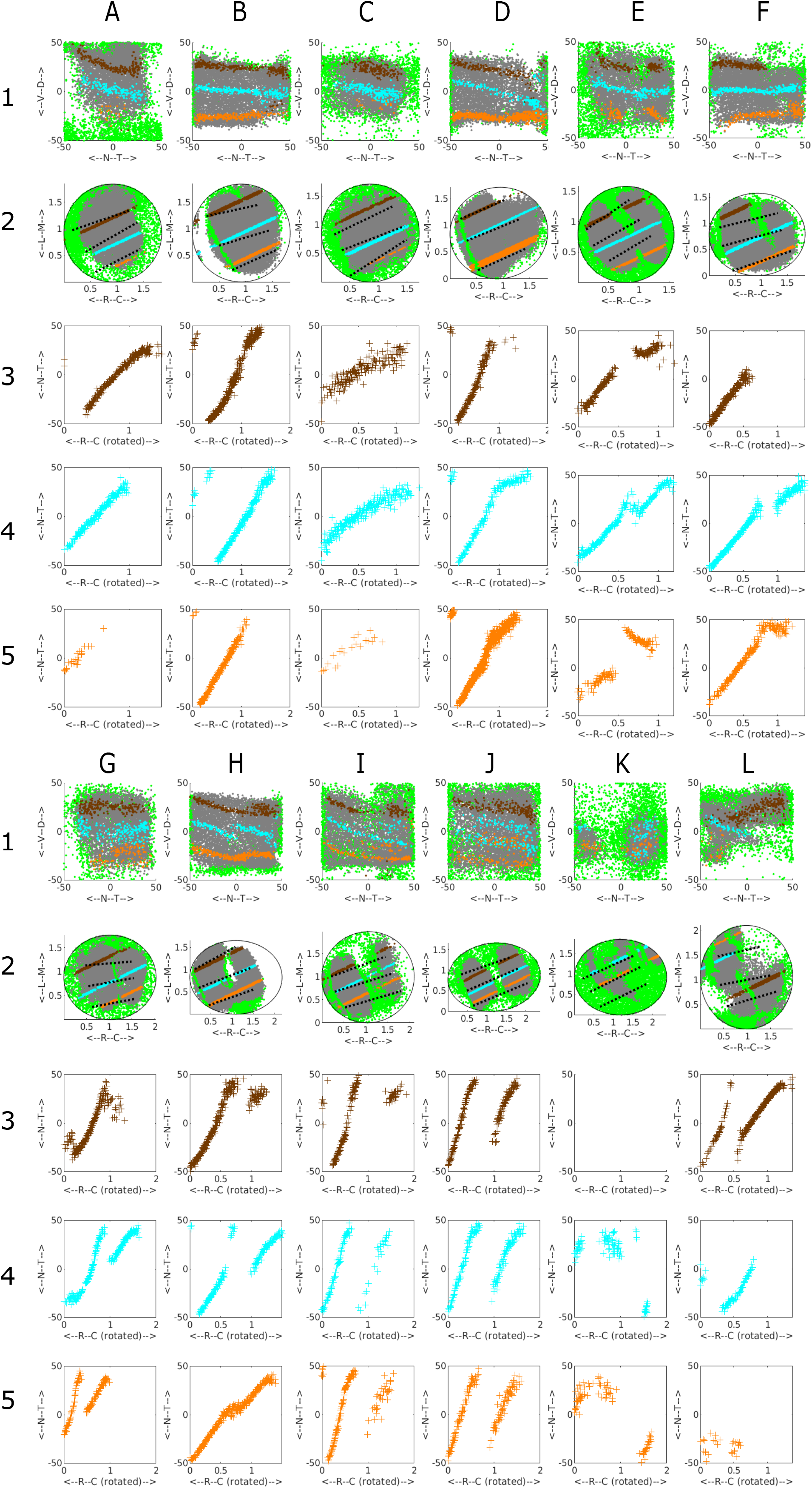
Examples from the database of the distribution of the variation of the azimuthal (nasotemporal) component of phase representing visual field position recorded along three straight line tracks drawn on the colliculus representing 1D sections at three standard elevations of -25° (orange), 0° (cyan) and +25° (brown) for WT, heterozygous (EphA3^ki/+^) and homozygous (EphA3^ki/ki^) knockins. **A, B**. WT. **C-H, K**. EphA3^ki/+^. **I, J, L**. EphA3^ki/ki^. Rows 1,2 show the distributions of eligible (grey) and rejected (green) pixels on the colliculus (row 2) and corresponding visual field positions (row 1). Each coloured line on the colliculus (row 2) is the straight line fit to the positions of all eligible pixels for which the corresponding visual field positions are at the given elevation. The coloured points plotted on the visual field representation is the projection back into the visual field from all the pixels on given straight line. These projections are not aligned perfectly with the appropriate elevations, suggesting that the nasotemporal and dorsoventral aspects of the map cannot be separated easily. Black dotted lines on the colliculus indicate the tracks used by Owens et al. (2015) in sampling the projection. Row 3-5: Plots of the azimuthal component of phase against collicular position along the appropriate track. The orientation of the 1D sections vary but all are seen to be at approximately 20° to the rostro-caudal axis. All three plots for the same dataset are drawn with the same horizontal scale, which varies from dataset to dataset. No data was recorded in the plot in Figure 2K, 3^rd^ row. Units of phase are degrees and those of position on the colliculus are mm.

In WTs (Figures 2A, 2B) there is one large area of eligible pixels on the colliculus (row 2), taken to be the main projection area, surrounded by rejected pixels. In Figure 2B there is a small and thin secondary projection area, seen also in Figure 2D. Heterozygous EphA3 knockins (HET; Figures 2C-2H) show a variety of projection areas. In some cases, there is one main area of eligible pixels (Figures 2C, 2D); in others there are signs of two substantial projection areas with rejected pixels between them of various forms (Figures 2E-2H). Homozygous EphA3 knockins (HOM), show a clear separation of the colliculus into two projection areas (Figures 2I, 2J). Figures 2K, 2L are examples of the three cases which did not show well-defined projection areas and so were not analysed further.

### 1D analysis: sampling the nasotemporal to rostrocaudal projection

To motivate our 2D analysis and compare our 1D results with that of Owens et al. (2015), in Figure 2 we also show the 1D projections at three standard elevations in the visual field (rows 3-5). The projections are from colliculus (row 2) to visual field (row 1). Most of the 21 cases analysed extensively each had a strong 1D map, extending from nasal field to cover most of the visual field (rows 3-5). In some of the profiles the map was absent or did not cover all of visual field (Figures 2A, 2C, row 5; Figure 2F, row 3), or had breaks (Figure 2E, rows 3,5; Figures 2F, 2H, row 4) In both the HOMs and the HETs, there was also a second projection from caudal colliculus of varying sizes, which was restricted to temporal field (Figures 2G-2J, rows 3-5 except Figure 2H, row 5). There were also other types of projection such as the small projection to temporal field from rostral colliculus (Figures 2B, 2D, rows 3-5).

In summary, the 1D projections are highly variable. Viewing how the tracks along which the 1D profiles were taken run over the irregular regions of eligible pixels on the colliculus (row 2) suggests that the 1D profiles reconstructed are sensitive to the exact positioning of the tracks. We conclude that assessing the projection at three standard positions does not allow for the variability in the maps and the nature of the projections is more complex than a binary classification into single or double projections (Owens et al., 2015). This gave additional motivation to pursue the construction of 2D maps.

### Wildtype 2D maps

To establish a baseline, the five WTs in the dataset were analysed. In each case there is a single large and extremely well-ordered projection, including two with a small aberrant projection.

Figure 3A shows the complete projection between each of the nodes on the colliculus (bottom) to the matching nodes in the visual field (top) for the WT shown in Figure 2A. The red links, mainly in temporodorsal visual field and caudomedial colliculus, indicate regions where links in the visual field cross, indicating violation of neighbourhood relations. Removal of selected nodes caused these links to be deleted until the largest ordered submap with perfect local order remained (Figure 3B). To give a sense of the overall polarity of the map and to relate these maps to the 1D slices shown in Figure 2A, three lines running nasotemporally which link the visual field nodes closest to the same standard elevations used in the 1D analysis were projected onto the colliculus. The map in Figure 3B has high local order with Map Quality, the percentage number of nodes in the largest ordered submap, of 96%. The rostrocaudal (RC) and mediolateral (ML) polarities, defined in Methods, are in excess of 90%.

**Figure 3.**
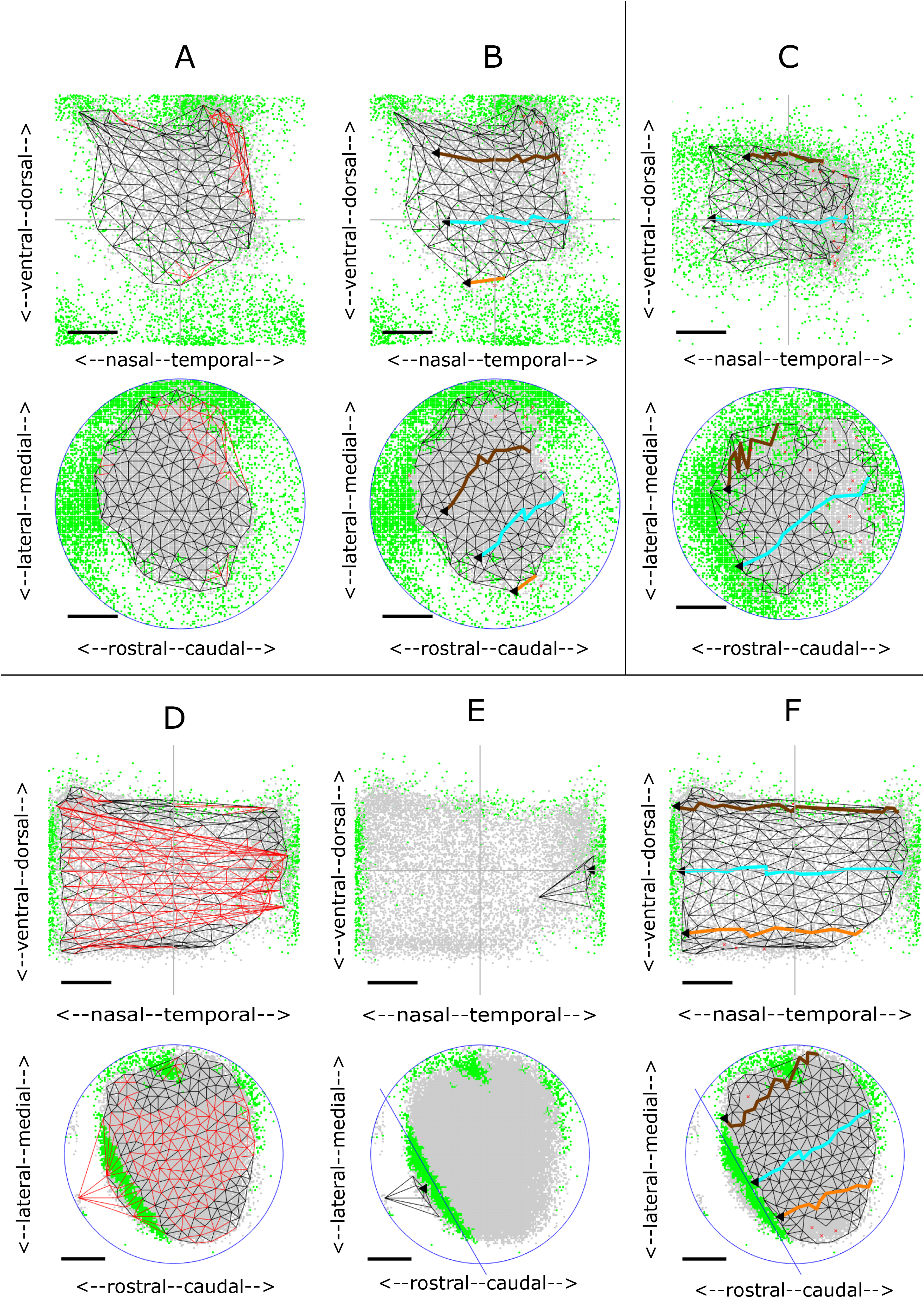
2D maps from colliculus (bottom) to visual field (top) for WT and HETA datasets. Neighbouring nodes on the colliculus and the corresponding nodes in the visual field map are connected by straight lines. Lines in the visual field which cross and the corresponding lines on the colliculus are coloured red. To relate the 2D maps to the patterns of 1D profiles in Figure 2, in some figures three lines representing the projections at -25° (orange), 0° (cyan) and +25° (brown) are shown. **A, B**. Maps formed for the WT shown in Figure 2A. **A**. The complete projection with 162 nodes. **B**. The largest ordered submap, constructed by removing 4% of the nodes from the complete projection to eliminate all crossing lines giving a Map Quality of 96%. RC, ML polarities: all 90%. **C**. The largest ordered submap for the HETA shown in Figure 2C, with 215 nodes in the whole map and 183 in the largest ordered submap giving a Map Quality of 86%. RC, ML polarities: all 90%. Azimuthal and Elevational Magnifications: 77°/mm, 60°/mm. **D-F.** Maps formed for the WT shown in Figure 2B. **D**. The complete projection, with 165 nodes. **E**, **F**. The two largest ordered sub-partmaps constructed on the two sets of nodes by partitioning the set of collicular nodes with a line drawn through the narrow strip of rejected pixels in rostral colliculus. **E**. Sub-partmap on rostral colliculus. **F**. Sub-partmap on the rest of the colliculus. RC, ML polarities are 94%. AM, EM: 115°/mm, 58°/mm. Scale bars: 20° (visual field); 200µm (colliculus).

Figures 3D-3F show another WT map with a strip of rejected pixels separating a large area of eligible pixels from a small area (Figure 2B). In the whole map (Figure 3D) there are many violations of neighbour relations. By dividing the collicular nodes into two by a line drawn through the strip of green rejected pixels and then constructing separate maps on the two collicular lattices so formed (Figures 3E, 3F), the small projection area rostrally was found to be associated with temporal field (Figure 3E). The massive violation of neighbour relationships seen in Figure 3D is due to this small part of temporal field being represented both rostrally and caudally.

In the group of five WTs, the number of nodes on the colliculus is 188±38, spanning an approximately 1 mm^2^ circular region. With a Map Quality of 91±4%, around 90% of the nodes are positioned in the correct relative order at an internode spacing of 80±8µm in the colliculus. The RC and ML polarities are 90±4% and 91±3%. The map is slightly more elongated along the (rotated) rostrocaudal axis than the (rotated) mediolateral axis. Azimuthal (AM) and Elevational magnifications (EM) are 93±13°/mm, 71±9°/mm.

### EphA3^ki/+^ 2D maps (HETS)

Two datasets had been discarded at the 1D analysis stage, one of these shown in Figure 2K. The remaining 14 cases fall into three groups.

#### HETA (5 cases)

In this group there is one projection area on the colliculus and a single ordered map. The largest ordered submap of one example is shown in Figure 3C, with pixel distributions in Figure 2C. These five maps are slightly less well-ordered than WTs, with Map Quality 86±2% against 91±4%. The number of nodes (190±22 against 188±38), the RC and ML polarity scores (91±2%; 90±1% against 91±4%; 91±3%) and the magnifications (98±16°/mm, 63±5°/mm against 93±13 °/mm, 71±9°/mm) are very similar to those in WTs (Figures 9A-9F).

In three cases, a small part of nasal field is associated with far caudal colliculus, one example of the pixel distribution shown in Figure 2D. The maps are similar to those found in two WTs (Figure 3D-3F). These suggest a slight phase miscalibration in the scanning. More generally, it demonstrates the importance of analysing the data appropriately.

#### HETB (4 cases)

The two regions of eligible (grey) pixels on the colliculus suggest two incompletely separated projection areas with rejected (green) pixels interposed (Figures 2E,2F). These could represent noise or regions associated with two distinct areas of visual field and thus cannot be mapped. Figure 4 shows the maps for these two HETs. Here, the rostral and caudal projection areas were defined by drawing a line through the narrow neck of the composite region to produce separate largest ordered sub-partmaps (Figures 4B-4C, 4E-4F). Map Quality computed over the two sub-partmaps is 3% more than in the single map.

**Figure 4.**
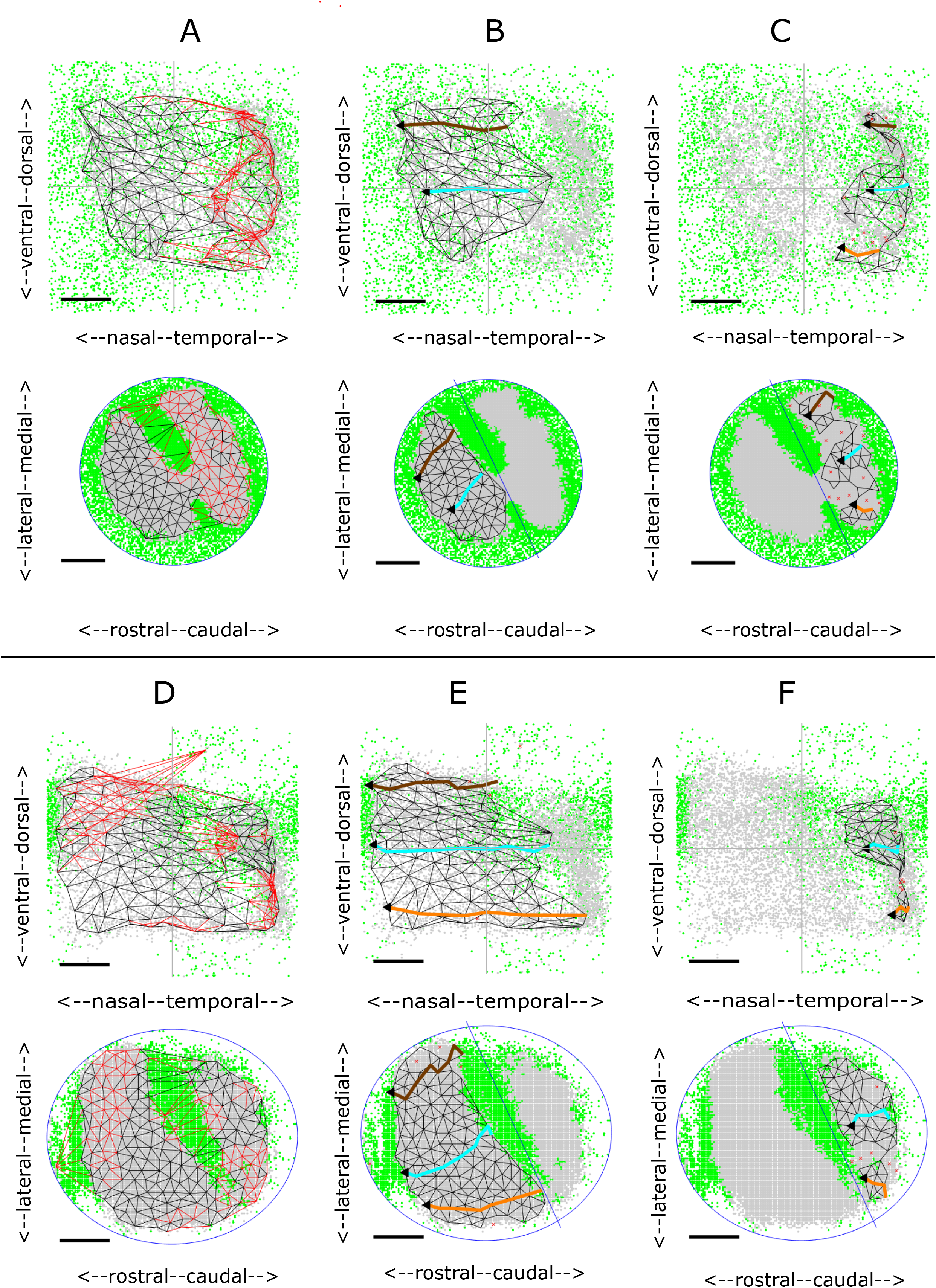
2D maps for two HETB datasets. **A-C.** Maps formed from the data shown in Figure 2E. **A.** Whole map with 148 nodes. Largest ordered submap has 124 nodes with Map Quality 84%. **B**, **C**. Sub-partmaps formed after partitioning the colliculus through the green areas of discarded pixels into rostral (**B**) and caudal (**C**) areas. 3% more nodes were recruited in the two sub-partmaps than in the largest ordered submap, giving a Map Quality of 87%. VFD, the area of visual field contained in both rostral and caudal projections: 3%. RC, ML polarities and AM, EM: 94%,89%, 118°/mm, 89°/mm (**B**); 68%, 83%, 84°/mm, 75°/mm (**C**). **D-F.** Maps formed from the data shown in Figure 2F. **D.** Whole map with 186 nodes. The largest ordered submap has 169 nodes giving a Map Quality of 91% **E**, **F**. Rostral and caudal sub-partmaps, with 3% more nodes than in the largest ordered submap. VFD: 7%. RC, ML polarities and AM, EM: 96%, 92% 101°/mm, 92°/mm (**D**); 68%, 92%, 86°/mm, 62°/mm (**E**). Conventions as in Figure 3.

In this category, the two separate partmaps make up a single map with only a very small area of visual field represented on both rostral and caudal subdivisions of the colliculus, the extent measured by the VFD (Visual Field Duplication). The area of visual field associated with rostral colliculus is 74±17% of the total occupied by the largest ordered submap against 30±15% to caudal colliculus, giving a VFD of 4±3%. Defining the total collicular region as the areas occupied by the rostral and caudal regions together with the area between them, the rostral region occupies 48±13% of the colliculus, the caudal region 33±10% and the central region 19±10%.

Map Quality of the two partmaps is 93±4% compared with 91±5% for the whole map, identical to the figure for WTs. The RC and ML polarities in both the largest ordered single sub-partmap and the largest ordered sub-partmap in rostral colliculus are all in excess of 90%, at the WT level, whereas for the smaller projection to caudal colliculus the polarities are slightly lower (Figures 9A-9D).

#### HETC (5 cases)

Here there are also suggestions of two projection areas (Figures 2G, 2H). In this group two well-ordered partmaps could be identified with polarity scores and Map Quality greater than for the whole map (Figures 9A, 9C, 9D). 91±11% of the visual field is represented in rostral colliculus and 27±14% in caudal colliculus giving larger VFDs than for HETBs, of 18±7%. Two HETCs, corresponding to Figures 2G, 2H, are shown in Figure 5. In both cases, whereas the projection from the rostral area is to the entire visual field (Figures 5B, 5E), there is no projection to nasal visual field from the caudal area (Figures 5C, 5F). This results in a higher AM than EM and significant VFD of 27% and 24% respectively (Figures 9E, 9F, 9B). The averages sizes of the rostral region of projection, the caudal region and the central region are similar.

**Figure 5.**
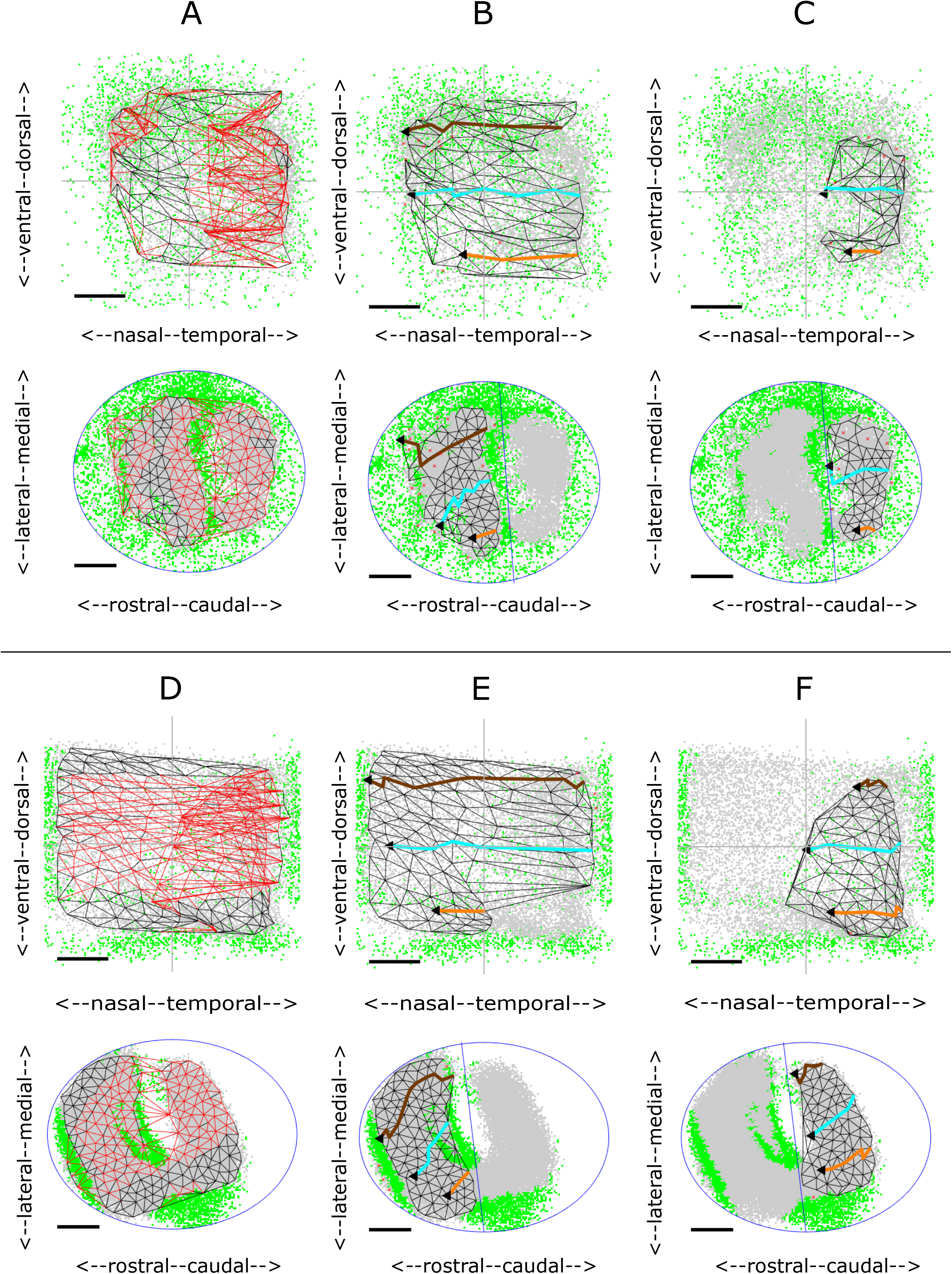
2D maps for two HETC datasets. **A-C.** Maps formed from the data shown in Figure 2G. **A**. Whole map with 180 nodes in the complete map. Largest ordered submap has 135 nodes with Map Quality 75%. **B**, **C**. Sub-partmaps, with Map Quality of 85% 10% more than in the largest ordered submap. VFD: 29%. RC, ML polarities and AM, EM: 91%, 85%, 168°/mm, 84°/mm (**B**); 90%,88%, 88°/mm, 73°/mm (**C**). The value of AM for **B** is almost twice that for **C**, primarily due to the high magnification in temporal field. **D-F.** Maps formed from the data shown in Figure 2H. **D**. Whole map with 207 nodes. Largest ordered submap has 188 nodes with Map Quality 91%. **E,F**. Sub-partmaps, with Map Quality 98%, 7% more than in the largest ordered submap. VFD: 29%. All polarities between 90% and 93%. AM, EM: 160°/mm, 82°/mm (**E);** 86°/mm, 76°/mm (**F).** Conventions as in Figure 3.

### EphA3^ki/ki^ (HOM) 2D maps

Four of the five available data sets were analysed, the fifth being rejected as a result of the 1D analysis (Figure 2L). In these four HOMs, partitioning the colliculus into two yields two distinct, well ordered partmaps. They have high polarity scores and Map Quality is 89±10%, substantially more than 71±10% for the whole projection (Figures 9A, 9C, 9D). The rostral projection area has a projection to almost all (98±4%) of the visual field whereas the caudal region has a projection to about one third (35±20%), in temporal field. As a result, the VFD is substantial, varying widely with individual scores of 17%, 25%, 29% and 59% of the visual field, an average of 33±18*%.* The sizes of the rostral projection area, the caudal projection area and the area between them are very similar (32±10%, 32±13%. 35±7%). The fact that a larger area of the visual field is associated with the rostral area than with the caudal area of the same size is reflected in the fact that the azimuthal magnification (AM) for the rostral partmap is double that for the caudal map: 203±20°/mm against 101±16°/mm, which is similar to the WT value of 93±13°/mm (Figures 9E,9F).

Figures 6A-6C show the HOM map with the smallest VFD, of 17%, with pixel distributions in Figure 2I. The projection from rostral colliculus covers most of the visual field (Figure 6B) and the small projection from caudal colliculus (Figure 6C) is to temporal field only. Figures 6D-6F show the HOM map with largest VFD, 59%, with pixel distributions in Figure 2J. Apart from far-nasal visual field, there is a double representation on the colliculus.

**Figure 6.**
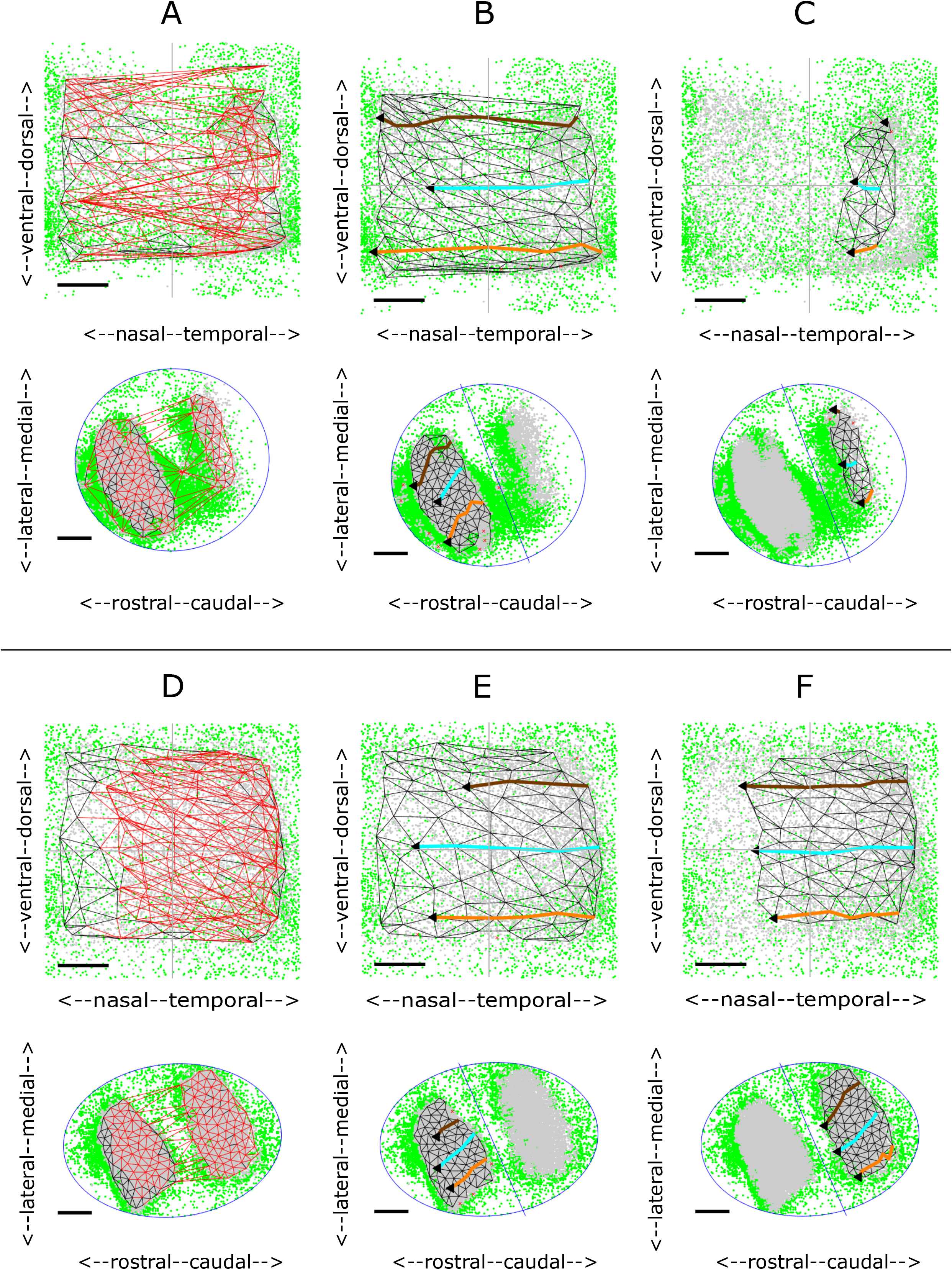
2D maps for two HOM datasets. **A-C.** Maps formed from the data shown in Figure 2I. **A**. Whole map with 148 nodes. Largest ordered submap has 123 nodes with Map Quality 83%. **B**, **C**. Sub-partmaps, with Map Quality of 93%, 10% more than in the largest ordered submap. VFD: 17%. All polarities between 86% and 91%. AM, EM: 212°/mm, 85°/mm (**B);** 79°/mm, 85°/mm (**C**). **D-F.** Maps formed from the data shown in Figure 2J. **D**. Whole map with 177 nodes. Largest ordered submap has 131 nodes with Map Quality 74%. **E**, **F**. Sub-partmaps, with Map Quality of 98%. 24% more than in the largest ordered submap. VFD: 63%. All polarities between 91% and 93%. AM, EM: 174°/mm, 117°/mm (**E); both**140°/mm, 85°/mm (**F).** Conventions as in Figure 3. Note that the homozygote map in Figures 6A-6C are from the data presented by Owens et al. (2015) as the paradigm heterozygote with a double map (their Figure 1N).

### Summary of 2D maps

Highly ordered 2D maps from colliculus to visual field were reconstructed from noisy Fourier imaging data derived from both WT and EphA3 knockin mice. Sampling the colliculus at a spacing of 80µm enables 2D maps to be constructed with around 90% of neighbouring points in the correct relative order and orientation.

In a WT map there is a single well-ordered map and HETA maps are identical to WT maps. In HETBs, HETCs and HOMs, the colliculus can be divided into two areas to produce two ordered maps. Figure 9A justifies this division, showing that more nodes are used in the two largest ordered partmaps than in the single map. The VFD in the two partmaps varies both within and between groups. On average it is greater in the HOMs, less in the HETCs, and very small in the HETBs (Figure 9B). In HETCs and HOMs, the area of visual field that is represented in both areas is restricted to temporal field, nasal field tending to be absent from caudal colliculus (Figures 5,6). Azimuthal magnification is significantly greater in the rostral map than in the caudal map, being greatest in the HOM maps and least in HETB maps (Figures 9E, 9F). Figures 9A-9F give summary statistics for all 21 cases.

The large degree of variability within both HET and HOM maps suggests that they are derived from a continuum of maps ranging from a single ordered map covering the colliculus to a map with a projection of visual field from two distinct areas on the colliculus. More of the visual field is represented rostrally than caudally where the representation of nasal field tends to be absent.

Since these maps are from colliculus to visual field, they do not give information about the individual projections of the EphA3+ and EphA3− cell types. It seems reasonable to conclude that in groups HETA and HETB, the projections from the populations of EphA3+ and EphA3− cells overlap completely or almost completely to give a single projection with little or no overlap in visual field, yielding effectively single wildtype maps.

In double maps, one possibility is that the rostral projection is from EphA3+ cells exclusively and the caudal projection from EphA3− cells exclusively. However, this implies that a substantial part of the representation of EphA3− cells on the caudal projection area would be missing (Figures 5C, 5F, 6C, 6F). It seems unlikely that these missing representations of nasal field are to the central part of the colliculus which cannot be mapped by Fourier imaging due to interference between two projections. This is because the two interfering projections would be from far-nasal and far-temporal field (green areas, Figure 1B), whereas in the HETC and HOM maps temporal field is always represented. Another possible explanation is that in temporal retina (corresponding to nasal visual field) there are no functioning EphA3− cells.

### Insights from modelling studies

To explore these issues requires study of double maps in which the projections from retina to the colliculus can be related to the projections from colliculus to visual field (or retina) supplied by Fourier-based intrinsic imaging. This information is available in modelling studies and we looked at double maps obtained from the energy minimisation model (Triplett et al., 2011). To compare with the experimental maps, we made Lattice plots of the maps from colliculus to retina after applying our simulation of the Fourier imaging process to the model data. These maps were supplemented by Lattice plots from retina to colliculus, including separate maps of the EphA3+ and the EphA3− projections.

Figure 7 shows a sequence of maps for different values of DR, the extra amount of EphA added to EphA3+ cells. With DR set at zero, in the collicular to retina projection, coloured black, rostral colliculus maps to temporal retina (visual field) and caudal colliculus to nasal retina (temporal field); there is a single map (Figure 7A). At a higher value of DR (Figure 7B), there is an area on the colliculus, coloured green and rejected by filtering because the average retinal position varies considerably. This defines two distinct projection areas. The projection from rostral colliculus covers most of the retina whilst that from caudal colliculus remains to nasal retina, giving a measurable VFD. As DR is further increased, the projection to nasal retina increases (Figures 7C, 7D), with larger VFD, until both rostral and caudal areas project to almost all of the retina (Figure 7E). Figures 9G, 9H shows statistics calculated over repeated runs on a smaller sized model to explore how Visual Field Duplication and average Map Quality depend on the value of DR.

**Figure 7.**
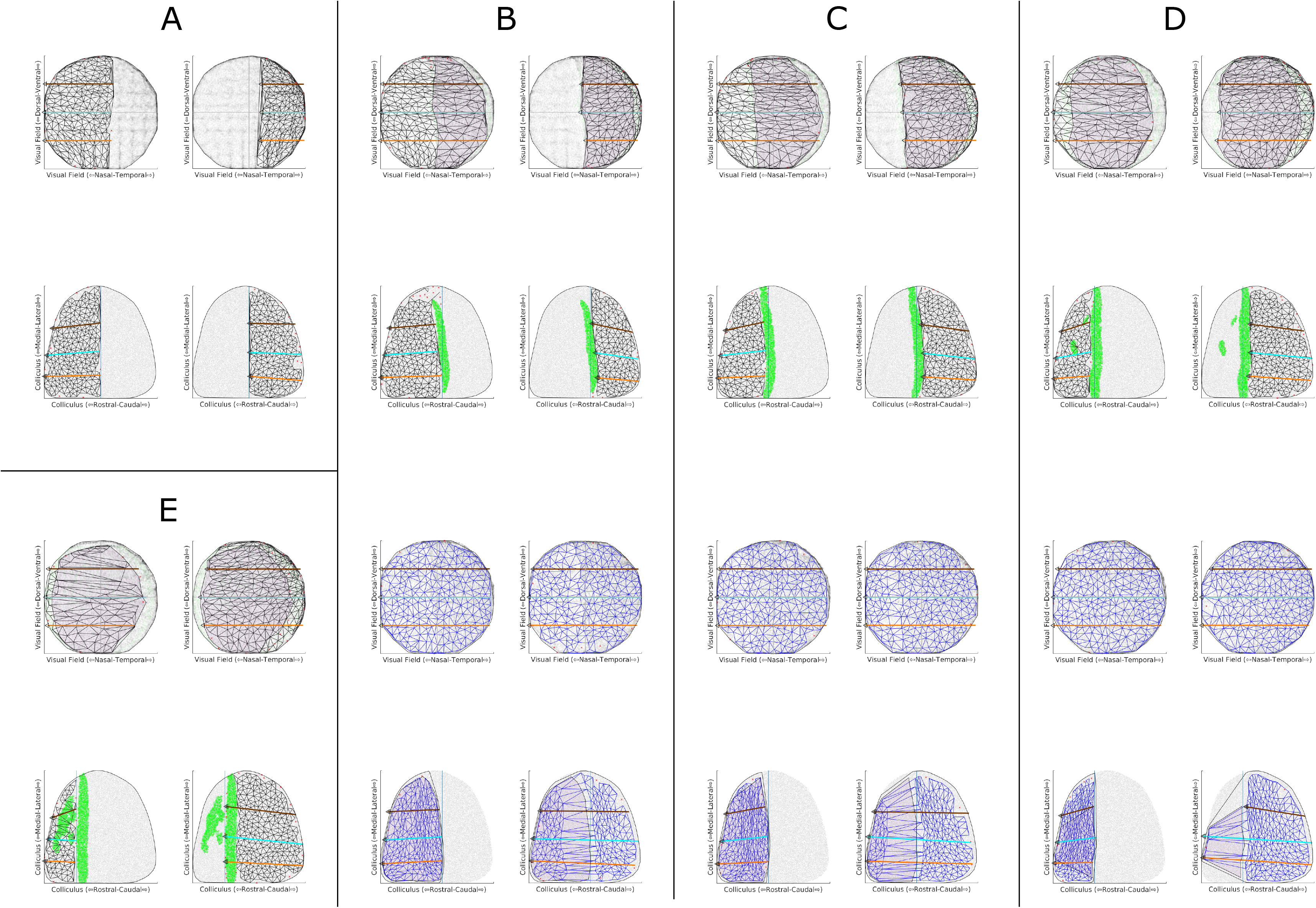
Single runs of the TK2011 model (Triplett et al., 2011). Connections developed between 10000 retinal cells and 10000 collicular cells, using the code due to Hjorth et al. (2015). Maps were computed for different values of DR, the amount of EphA added to the Islet-2 cells, and constructed using the Lattice Method (Willshaw et al., 2014). Largest ordered sub-partmaps are shown. Retina is plotted as visual field to aid comparison with the experimental plots. For the mapping from colliculus to retina, coloured black, the collicular lattice was divided into two by a line through the centre of the green (rejected) pixels. The lefthand plot is the map from the rostral part of the colliculus; the righthand plot that from the caudal part. For the mapping from retina to colliculus, coloured blue, the lefthand plot is the projection from EphA3+ cells and the righthand plot that for the EphA3− cells. Map orientation shown as in the experimental plots. **A.** DR=0. Colliculus to retina map only. The collicular lattice was divided into equal-sized rostral and caudal halves. **B-D.** Projections in both directions are shown for DR = 0.22, 0.34, 0.56. For the projection from colliculus to retina the shaded region in the retina is represented in both rostral and caudal collicular areas. **E**. DR=1.12. Colliculus to retina projection only. Each retinal cell was chosen to be EphA3+ with probability 0.5. Parameter values. α=90, β=135, a=0.03,b=0.11 (Triplett el., 2012; Hjorth et al., 2015); γ=0.00625, 5X10^8^ iterations, (Gale, 2022). The EphA gradient is derived from Reber et al. (2004). Values of DR quoted are expressed as a fraction of the span of the values of EphA over the WT retina. On this scale, the values of DR for an EphA3^ki/+^ and an EphA3^ki/ki^ map (Reber et al., 2004) are 0.37 and 0.74 respectively.

| Figure | A | B | C | D | E |
| --- | --- | --- | --- | --- | --- |
| DR | 0.0 | 0.22 | 0.34 | 0.56 | 1.12 |
| Number of nodes | 485 | 436 | 415 | 426 | 407 |
| VFD | 1 | 47 | 58 | 74 | 76 |
| MAP QUALITY (largest ordered submap) | 97 | 80 | 74 | 54 | 73 |
| MAP QUALITY (largest ordered partmaps) | 98 | 95 | 96 | 95 | 95 |

These simulated maps from colliculus to retina can be related directly to the experimental maps and have several similar features. Simulated WT maps are single. A simulated EphA3 knockin map can be single, or a double map where the proportion of the retina (visual field) represented in both projection areas varies widely, depending on the value of the EphA3 augment, DR. In the double maps a larger extent of the retina is represented rostrally than caudally with much of temporal retina (nasal field) not represented in the caudal area. In the experimental maps there is significant variation, such as within the HOM category where either a small or a large extent of visual field is represented in rostral colliculus (Figures 5C, 5F). This difference could be attributed to the fact that, unlike in the simulations, signal strength varies considerably over the colliculus (Kalatsky and Stryker, 2003; Cang et al., 2008). The smaller amount of colliculus available for constructing the map after filtering may also explain why in the simulations, ordered maps of 400-500 nodes were constructed compared to 150-200 for the experimental maps.

Examination of the simulated projection from retina to colliculus now enables identification of the separate projections from the EphA3+ and the EphA3− cells. As described above, for DR=0, both EphA3+ and EphA3− cells from all parts of the retina project across the entire colliculus to give a single map. For a larger value of DR (blue maps, Figure 7B), the projection from EphA3+ cells is restricted to rostral colliculus and EphA3− cells project more widely so that rostral colliculus is innervated by both EphA3+ and EphA3− cells. At a larger value, the projection from EphA3− cells starts to become restricted to caudal colliculus with a strip of double innervation rostrally (Figure 7C), which becomes smaller for larger DR. (Figure 7D). Ultimately EphA3+ cells project rostrally and EphA3− caudally to give a double representation of retina on the colliculus. Figure 8 shows a set of 1D profiles for a series of maps with different values of DR, which also illustrates how these maps are formed, as described in the Legend. Rows 3 and 4 in Figure 8 illustrate the difference between a 1D map of the retina (visual field) seen from the colliculus and that of colliculus seen from the retina.

**Figure 8.**
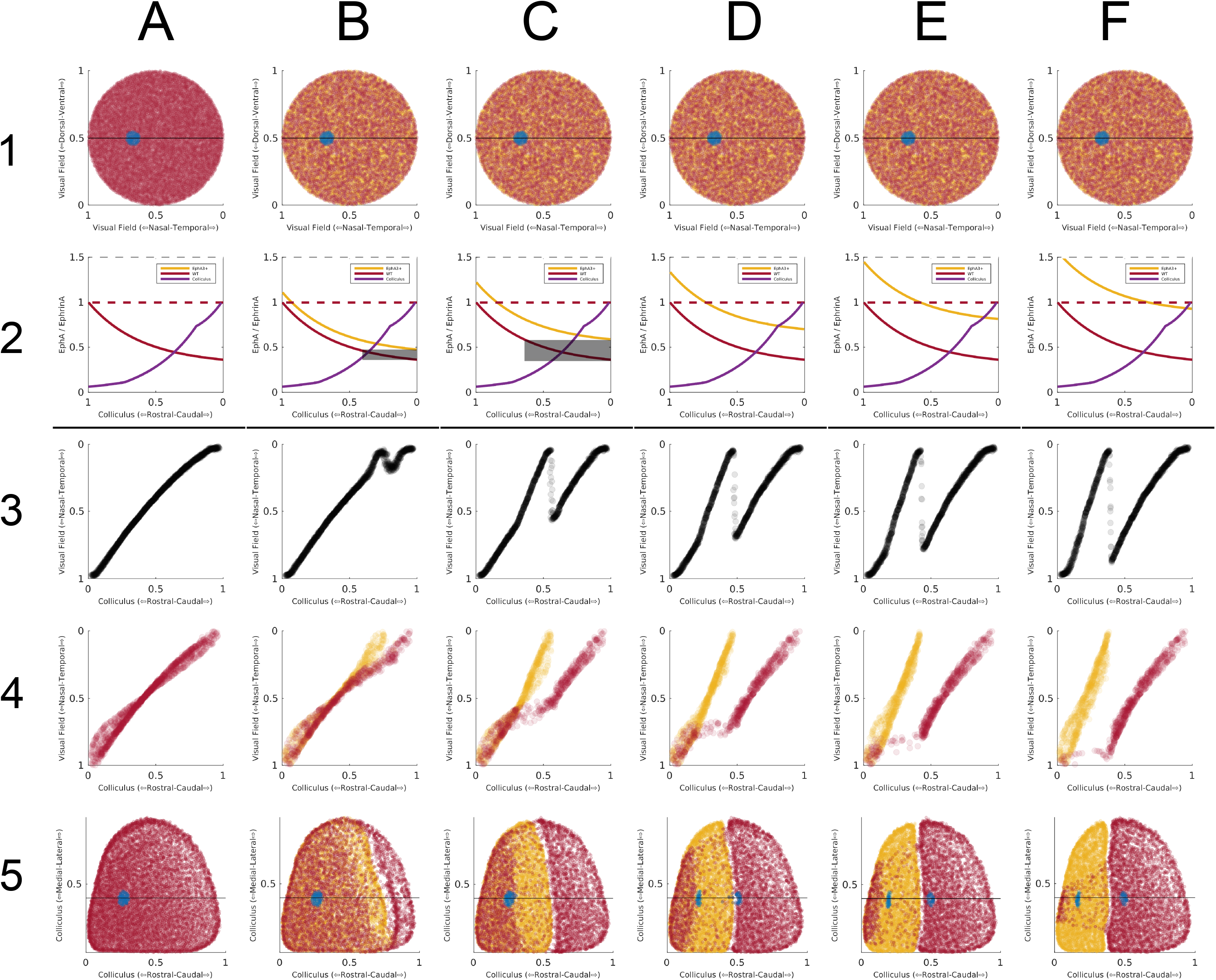
Data of the type used to generate Figure 7 from the TK2011 model was plotted to show a sequence of six 1D projections between the retina measured along a straight line running nasotemporally and the colliculus measured along a straight line running rostrocaudally, for increasing values of DR. Retina plotted as visual field. Each subfigure has 5 separate plots. Row 1: Distribution of EphA3− (purple) and EphA3+ (orange) cells over the retina. The horizontal line indicates the line along which the 1D profile was taken. Blue marks the site of a simulated injection of 1% of the retinal cells (row 1) and its projection on the colliculus (row 5). Row 2: The distribution of EphA along the nasotemporal axis of the retina for both EphA3− cells (purple) and EphA3+ cells (orange) and of ephrinA cells (magenta) across the rostrocaudal axis of the colliculus. Row 3: Nasotemporal component of the positions of all retinal cells with contacts from the colliculus cells lying along the rostrocaudal axis of the colliculus at mediolateral position 0.5, marked in row 5. Row 4: Rostrocaudal component of the positions of all collicular cells with contacts from the retinal cells lying along the nasotemporal axis of the retina at dorsoventral position 0.5, shown in row 5. The projections from the EphA3− cells (purple) and the EphA3+ cells (orange) can be distinguished. Row 5. Distribution of the average positions on the colliculus of the projections from each retinal cell, with the projections from the injected cells in blue. Initially a large number of contacts are made at random and then contacts are withdrawn selectively, those that lower the overall energy being maintained. The difference in energy between the contacts made by two retinal cells on the same collicular cell is the product of the amounts of EphA and ephrinA present. The cell with higher energy tends to be withdrawn as this decreases the energy by a larger amount. Note that, as in all Markov Chain Monte Carlo methods (Brooks et al., 2011), this is not a true minimisation model which converges to a single configuration, but rather to a distribution of possible configurations (Gale, 2022). **A.** DR=0.0. EphA3− and EphA3+ cells at the same retinal location have the same amount of EphA and so there is a single map (Figure 7A). **B.** DR=0.16. Some WT cells from nasal retina (temporal visual field) have values of EphA lower than those in the EphA3+ population at the same collicular position (row 2, grey bar) and so survive at the expense of the contacts from the EphA3+ cells there. The contacts made more centrally by these same EphA3+ cells now compete against the EphA3− cells there, which have higher energy. These EphA3+ cells tend to survive, and the overall effect is that the EphA3+ projection from nasal retina is more central. **C.** DR=0.32 (Figure 7B); **D.** DR=0.48 (Figure 7C). There is a growing population of EphA3− cells with unique values of EphA, increasing the extent of the single EphA3− projection caudally and leading to contacts from more and more EphA3+ cells being removed from this region of the colliculus. There is still a double projection far rostrally, where the activity mechanism favouring neighbour-neighbour connections dominates over the weaker chemoaffinity mechanism. **E**. DR=0.64; **F**. DR=0.78 (Figure 7D). A very strong EphA3+ projection centrally, results in increased pressure to maintain neighbourhood relations there by lowering the energy of the EphA3− contacts which out-compete the EphA3+ contacts and effectively break the EphA3− projection there. As DR is increased, the effect of chemoaffinity rostrally becomes stronger, diminishing the superposed projection. For sufficiently large DR, all the EphA3− cells have values of EphA lower than the EphA3+ cells and a double projection results. EphA3+ cells project rostrally and EphA3− cells caudally.

There are two other relevant properties of the model (Gale and Willshaw, in preparation).

1. The level of neural activity controls the tendency from neighbouring retinal cells to connect to neighbouring collicular cells. Reducing activity slows down the development of the projection in its earliest stages. This also reduces the number of EphA3+ and EphA3− cells from the same retinal location projecting in register on rostral colliculus.
2. In double maps the distribution and relative proportion of EphA3+ to EphA3− cells in the retina affect their patterns of innervation. For example, changing the percentage of cells that are EphA3+ changes the proportions of the colliculus innervated by the two populations.

In these simulations it is how the profile of EphA over the EphA3− cells compares with that over the EphA3+ cells which is crucial in determining the form of an EphA3 knockin map. As seen by Fourier-based imaging, if the two profiles are almost identical, a single or near single map would result (HETA, Figure 3; HETB: Figure 4; Figures 7B, 7C). If the two profiles are distinct, a partially double map results (HETC. Figure 5; HOM: Figure 6). According to the characterisation of the EphA gradients by Reber et al, (2004) used in the model, Figures 7B and 8C would represent an EphA3^ki/+^ map and Figures 7D and 8F an EphA3^ki/ki^ map.

We repeated the simulations of the computational model which Owens et al. (2015) used to support their stochastic hypothesis. This model, TK2006 (Tsigankov and Koulakov, 2006, 2010) is an earlier version of the one that we used in our analysis of the experimental data. We use this model to give a direct comparison; we did not use this model in our analysis of the experimental data as it allows only one contact per cell, which disrupts the mapping, as described below.

We used the same set of parameters as Owens et al. (2015) and varied the magnitude of the EphA3 augment, DR, over the same range. The maps in the first row of panels of Figures 9I-9M resemble their published maps. At low DR, there are single maps (Figure 9I) and at higher values completely double maps (Figure 9M). With a change of DR from 0.3 (Figure 9I) to 0.36 (Figure 9J), a single map became a blotchy map of the type that Owens et al. (2015) found, which transformed gradually into a double map over the range DR =0.36 to DR=0.7. (Figures 9J-9M). In contrast, Owens et al (2015) reported that a slight variation of DR around 0.36 would switch the map from single to double. Plotting the projections from the EphA3+ and EphA3− retinal cells separately (rows 2 and 3, Figures 9I-9M) revealed that the blotchy maps are due to neighbouring cells of the two types forming interleaving projections, each excluding the contacts from cells of the other type, similar to a pattern of zebra stripes This is a result of the strong competitive constraint of requiring one-to-one connectivity which masks the smooth variation from a single projection to a double projection as DR is increased.

**Figure 9.**
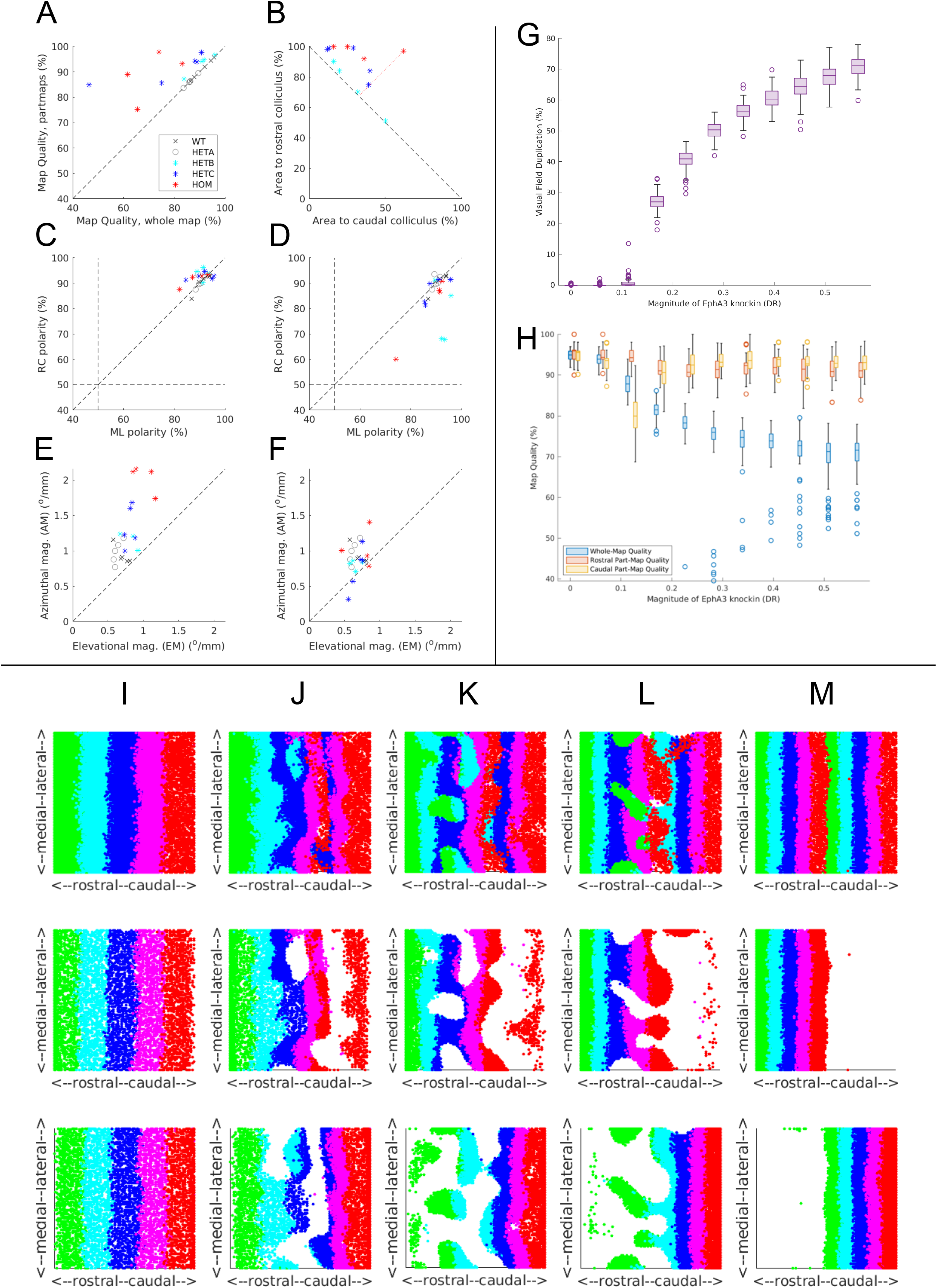
Summary and controls for both experimental and modelling data. **A-F.** Summary data for the 21 experimental maps analysed. **A**. Map Quality of the largest ordered two partmaps plotted against Map Quality for the largest ordered submap. For the single maps identified in the WT and HETA categories, the same Map Quality is plotted on both axes. Data points above the diagonal indicate that the partmaps cover more of the colliculus than the whole map. **B**. For the HETs and the HOMs where two projection areas were identified, the relation between the area of visual field represented in the rostral area compared to that in the caudal area. VFD is the perpendicular distance from the given point to the off-diagonal. **C**, **D.** Rostrocaudal (RC) polarity plotted against mediolateral (ML) polarity in double maps for the rostral map (**C**) and the caudal map (**D)**. **E**, **F.** Azimuthal magnification (AM) plotted against elevational magnification (EM) in double maps for the rostral map (**E**) and the caudal map (**F)**. In **C-F**, data from the single maps is included. **G, H.** Summary data for multiple simulations carried out on the TK2011 model of reduced size for different amounts of the EphA3 knockin, DR. **G.** VFD. **H**. Map quality for both largest ordered submaps and partmaps. 2000 retinal cells and 2000 collicular cells, 2X10^7^ iterations per run. 100 runs for each value of DR. Mean shown by the horizontal line within the box, which encloses one quarter of the data points. Standard deviation shown by the vertical line. **I-M.** Maps generated by simulations of the TK2006 model (Tsigankov and Koulakov, 2006) for five different values of the EphA3 augment, DR, to be compared with those in Figure 7 of Owens at al. (2015), using the same code (Reber, pers. comm.). The retinal cells are labelled according to their position along the nasotemporal axis by five stripes of different colours (Owens et al., 2015). For each value of DR, there are three maps showing the colour-coded nasotemporal origins of the retinal cells (plotted as visual field) projecting to each part of the colliculus. Row 1: projection from all cells; row 2: projection from EphA3+ cells; row 3: projection from EphA3− cells. **I**: DR=0.3 (0.35); **J**: DR=0.36 (0.42); **K**: DR=0.4 (0.46); **L**: DR=0.5 (0.58); **M**: DR=0.7 (0.81). In the TK 2006 and TK2011 models the values of DR were normalised to different bases. To aid comparison with the simulations shown in Figures 7-8, the corresponding values for TK2011 are shown in parentheses.

In summary, our simulations with this model shows that, as DR is increased from 0.3 to 0.7, the map evolves gradually from a single into a double map, by DR=0.7. We did not find any evidence for small variations of DR producing rapid switching between the single and double states.

## Discussion

In the original analysis of this dataset, Owens et al. (2015) sampled data from the azimuthal scan along three fixed rostrocaudally orientated tracks to produce 1D maps, defined as single or double using their quantitative measure. In different EphA3^ki/+^mice, at the three locations sampled they found single maps (as in WTs), double maps (as in EphA3^ki/ki^ mice) or a mixture of the two at different positions along the mediolateral axis (Figure 1A-1C). Eliminating both genetic variation and variation in neural activity, they concluded that the mixed category of map represents an unstable transitional state between those of a single and a double map, which can be interchanged by small fluctuations in the competing forces of chemoaffinity and neural activity acting on the innervating nerve fibres.

Our 2D maps were constructed from data from both elevation and azimuthal scans using the Lattice Method (Willshaw et al., 2014). This is a general method for constructing topographic maps. It provides quantitative measures of local and global ordering which can be used to compare maps without assuming a particular map scale or orientation. In situations where large datasets are available, statistics derived from this method can be used in population studies generated from experiment or from simulation (Figures 8G, 8H; Gale, 2022). Sampling the colliculus at a resolution of 80 µm enables 2D maps to be constructed in which over 90% of neighbouring points are in the correct relevant order and orientation. Applying the Lattice Method to a different dataset had given a spacing of 54µm (Willshaw et al., 2014); with standard intrinsic imaging it was 120 µm (Mrsic-Flogel et al., 2005) and for extracellular recording 200µm (Drager and Hubel, 1976). This method has wide applicability and can be applied to other datasets such as from mouse derived using ultrasound (Mace et al., 2018) and from Xenopus using Calcium imaging (Li et al., 2022).

In both EphA3^ki/+^ and EphA3^ki/ki^ mice we found a variety of maps, ranging from an entirely single projection to a double map with a large proportion of the visual field projecting to the rostral area and a smaller proportion, consistently from temporal visual field, to the caudal area. Owing to a lack of data we could not construct 2D maps for the small number of cases relating to the roles of genetic diversity and the neural activity that Owens et al. (2015) analysed. Their interpretation relies heavily on the measure they developed for classifying 1D maps as single or double which requires validation using ideal data.

The Fourier-based intrinsic imaging method provides a one-to-one mapping from colliculus to visual field. When applied to double maps, filtering is required to remove the ambiguous reports from the parts of the colliculus where distinct areas of visual field project together. The main difference between our approach and that of Owens et al. (2015) is that we constructed full 2D maps using data from both azimuthal and elevational scans rather than sampling the 1D projection from unfiltered data from the azimuthal scan

Owens et al. (2015) also presented results from a computational model showing the type of stochasticity that they advocated. However, when we reran their computer simulations using the same parameter values, we found no evidence for stochasticity. When we ran a more biologically realistic variant of the same model (Triplett et al., 2011; Hjorth et al., 2015), the results were broadly consistent with our maps of the experimental data. Although we could not analyse the maps found in EphA3 knockin animals also subject to knockout of the β2 component of the acetylcholine receptor, which affects neural activity (Pfeiffenberger et al., 2006; Cang et al., 2008), we found that lowering the strength of activity in the model reduces the extent to which EphA3+ and EphA3− cells project in register on rostral colliculus.

The chemoaffinity mechanism used in the model TK2011 (Triplett et al., 2011; Hjorth et al., 2015) which we analysed in detail is of the graded matching type (Prestige and Willshaw,1975). The model TK2011 fits the experimental data qualitatively. According to this model, the projections from EphA3+ and EphA3− retinal ganglion cells are superposed in rostral colliculus. As first pointed out by Reber et al. (2004) in presenting their relative signalling model, this superposition is a property of a graded matching model where in rostral colliculus there is a relatively small difference between the chemoaffinity signal carried by the two classes of innervating retinal fibres. The TK2011 model is not unique and has the disadvantage that the notion of energy minimisation is difficult to translate into the biology. Other simpler graded matching models, such as involving the gradual addition of contacts (Prestige and Willshaw, 1975) are feasible.

The consistent absence in our double maps of a projection from nasal field to caudal colliculus could be caused by EphA3− cells and EphA3+ cells of temporal origin projecting in register to rostral colliculus, in line with axonal tracing (Brown et al., 2004; Reber et al., 2004; Bevins et al., 2008; Owens et al. 2015) and as found in the modelling. This effect could be investigated in each particular case by looking for a coincidence of EphA3− and EphA3+ fibres projecting to the appropriate part of the colliculus. For example, in the case shown in Figure 6A-6C, there should be a large, superposed projection in medial colliculus, for Figures 6D-6F, there should be little or none. If there is no superposed projection, the missing part-projection, presumably from the EphA3− cells, must be made elsewhere. Alternatively, there could be an inhomogeneous retinal distribution of the Islet2+ cells to which EphA3 is attached. Pak et al. (2004) found a weaker representation of Islet2+ cells in the ventrotemporal crescent of the retina than elsewhere in the retina. Marcucci et al.(2018) found that neurogenesis of retinal ganglion cells in ventrotemporal retina lags behind that in dorsonasal retina.

### Alternative hypotheses

The hypotheses compatible with the original 1D analysis are: (1) in EphA3^ki/+^ maps there is stochasticity in the mechanisms making the connections on the colliculus (Owens et al., 2015); (2) the level of EphA3 in Islet2 cells varies systematically over the retina. Our 2D maps present a different picture of the data from the original 1D analysis which seems to rule out both hypotheses. In the EphA3 knockins we found a variety of double maps from both EphA3^ki/ki^and EphA3^ki/+^ animals where the variation is along the rostrocaudal axis, which concords with the anatomical evidence (Brown et al., 2000); we did not find any mixture of single and double maps at different locations along the mediolateral axis.

The substantial variability in the extent of the double projections from animal to animal occurring in both genotypes suggests another hypothesis: (3) the relative amount of the induced EphA3 receptor compared with the levels of endogenous retinal EphA varies between animals. A similar systematic variability is seen in the computer simulations as the level of EphA3 is increased from zero (Figure 7, Figure 8, Figure 9G). Several authors have quantified the amount of EphA3 in Islet-2 retinal ganglion cells. Brown et al. (2000) used in situ hybridisation and presented their data schematically. On the basis that mRNA measurements they inferred that the level of EphA3 in the homozygous knockin is twice that of the heterozygous knockins. Reber et al. (2004) showed that the levels of EphA3, EphA4, EphA5 and EphA6 in the retina have a standard deviation of around 10% for each receptor type. It is not clear whether these results are from one particular animal or averaged over different animals. A possible mechanism for a variation in the level of induced EphA3 between animals is haploinsufficiency. This is well known in Drosophila (Salazar and Yamamoto, 2018) and has been identified for EphA7 in humans (Levy et al., 2021) but there is no direct evidence for it in EphA3-Islet2 mice.

To explore the consequences of the third hypothesis requires examining how the two populations of EphA3− and EphA3+ retinal cells project to the colliculus and relating the form of each functional EphA3 knockin 2D map to (i) the distribution of Islet2 cells over the retina, the density of EphA3 in different heterozygotes and (ii) the location of collicular cells which are innervated by both EphA3+ and EphA3− cells. The Fourier-based imaging method has the advantage of providing data from over the entire brain structure assessed but it requires surgical intervention, making it difficult to combine with methods targeting one class of retinal ganglion cells. Methods using tangentially inserted high density electrodes (Sibille et al., 2022) have been used in mouse superior colliculus and provide 200 recording sites but along only one axis. Functional ultrasound may be more suitable, being applied externally. This method has been used already to provide phase maps in mouse colliculus (Mace et al., 2018). Combining functional ultrasound with optogenetics (Brunner et al., 2020) to look at the EphA3+ projection from the Islet2 cells alone may be possible to avoid the problem of ambiguous responses from part of the colliculus holding projections from widely different part of the visual field.

## Author Contributions

DJW secured funding, conceived the project, carried out the experimental data analysis and wrote the paper. NMG secured funding, collaborated in project design, carried out the computational modelling and wrote the paper.

## Acknowledgements.

This research was funded in whole, or in part, by the Wellcome Trust (Programme Grant number W083205A (DW) and PhD grant (number 215153/Z/18/Z (NG)). Our thanks for support to members of ANC (Edinburgh) and DAMTP (Cambridge) and to Matt Nolan, Stephen Eglen, David Price and Peter Dayan for their comments. Special thanks to Jason Triplett for sharing the experimental data with us and for his comments. For the purpose of open access, the authors have applied a CC BY public copyright licence to any Author Accepted Manuscript version arising from this submission.

